# Polystyrene Nanoplastics Disrupt Mouse Placenta Development in a Sex-Dependent Manner

**DOI:** 10.64898/2026.05.22.727211

**Authors:** Hanin Alahmadi, Allison Harbolic, Christopher De Oliveira-Cordova, Raulle Reynolds, Michelle Jojy, Courtney Potts, Sofia Doan, Tanvi Mathur, Mohammad Saiful Islam, Melisa J. Andrade, Quinton Smith, Phoebe A. Stapleton, Somenath Mitra, Genoa R. Warner

## Abstract

Plastic production has been increasing exponentially. Throughout their lifespan, plastics degrade into smaller particles that accumulate in our bodies and the environment. Recent studies found these plastic particles can cross the placental barrier and reach the fetus. However, the impact of plastic particles on placental function is still unknown. We hypothesized that nanoplastics would disrupt placental growth and function, specifically focusing on transforming growth factor beta (TGFβ) signaling. To understand the impact of plastic particles on the placenta, we orally exposed pregnant CD-1 mice to 50 nm or 200 nm polystyrene plastic particles from gestation day 8 to day 15 at a human-relevant concentration of 5 mg/kg/day. After euthanization on day 15, placenta and fetus weights were recorded, and tissues were prepared for histomorphology and gene expression analysis. We observed a statistically significant decrease in the area of the decidua in the placentas for the 200 nm treatment group and a borderline significant decrease in decidua area for the 50 nm treatment group compared to control. However, when we separated by sex, only the male decidua were significantly decreased in the 200 nm group. Gene expression analysis of key signaling factors in the TGFβ pathway identified increased expression of *Smad2* and *Smad3*, which may be suppressing prolactin and estrogen receptor signaling. Overall, both particle sizes disrupted placenta structure and signaling in a sex-dependent manner and may be acting as endocrine disruptors.

## 1 Introduction

Plastics have become integral to modern society due to their durability, low cost, and versatility, leading to a significant increase in global production over recent decades. In 2021, global plastic production was estimated at approximately 390 million metric tons, reflecting a substantial increase from previous decades, with projections indicating continued growth in the coming years [1]. This surge has resulted in widespread environmental accumulation of plastic waste, which, through degradation processes, breaks down into microplastics and nanoplastics (MNPs) [2]. These particles are now present across various ecosystems and have been detected in human tissues and fluids, raising concerns about their potential health impacts [3].

Emerging research indicates that nanoplastics can cross biological barriers, including the placenta, a critical organ responsible for nutrient and waste exchange between mother and fetus, as well as hormonal regulation during pregnancy [4]. The placenta plays a vital role in fetal development and serves as a selective barrier against environmental contaminants. However, recent studies have detected the presence of MNPs in human placental tissues [5,6]. In human term placentas, MNPs were measured in all tested samples at an average of 127 µg/g, with maximum levels of 685 µg/g [6]. Additionally, animal studies have demonstrated that maternal exposure to NPs can lead to their accumulation in both placental and fetal tissues, potentially resulting in adverse developmental outcomes [7,8]. Specifically, particles up to 240 nm have been shown to perfuse to the fetus in humans and up to 500 nm polystyrene spheres can accumulate in the placenta in mice [9,10]. Because of their small size, large surface area to volume ratio, and ability to translocate into cells, NPs are considered to be of greater toxicity risk compared to larger particles [11].

The infiltration of NPs into the placenta raises significant concerns regarding their potential to disrupt placental function and fetal development [12,13]. Recent human epidemiology has found associations between placental microplastics and decreased fetal size, indicating further study is warranted [14–16]. Impairment of placental development can lead to pregnancy complications such as fetal growth restriction, preeclampsia, pre-term birth, stillbirth, miscarriage, and maternal hemorrhage [17]. In rodents, NPs have been shown to alter placenta size, weight, and layer area [12,18–21], although specific effects and magnitude vary across particle properties and experimental conditions. In addition, NP exposure disrupts molecular signaling pathways such as PERK/eIF2α/ATF4 and hormone levels (estrogen, progesterone) [21]. In vitro, unmodified and carboxylated NPs migrated into and disrupted invasion and migration in trophoblast cells [22,23]. Although multiple studies report associations between NP exposure and altered placenta morphology, little is known about the mechanisms of action.

Transforming growth factor beta (TGFβ) signaling plays an essential role in early placentation as well as vascularization and angiogenesis in later pregnancy [24]. Dysregulation of TGFβ signaling is implicated in intrauterine growth restriction (IUGR), preeclampsia, and placenta accreta syndrome [24–26]. TGFβ also regulates estrogen signaling [27]. In an adult model, exposure to 500 nm nanoplastics increased TGFβ levels in rat ovaries, but this pathway has not been investigated in the placenta [28]. Thus, the objective of this study was to evaluate the impact of developmental nanoplastics exposure on TGFβ signaling in the placenta. We hypothesized that oral nanoplastic exposure in mice during the period of placenta formation would result in placental translocation of particles, altered placenta morphology, and changes in expression of genes related to placenta functions such as hormone synthesis, metabolic regulation, invasion, adhesion, and vascular growth and remodeling, including the TGFβ signaling family.

## 2 Method and Materials

### 2.1 Nanoplastics

1% suspensions of 200 nm red fluorescently labeled polystyrene and 50 nm blue fluorescently labeled polystyrene nanospheres were purchased from Fisher Scientific (B50 and R200, respectively). Red and blue nanoplastic particles were characterized by scanning electron microscopy (SEM) and dynamic light scattering (DLS) to confirm shape and size.

### 2.2 Animals and study design

Adult CD-1 mice at 60 days of age were purchased from Charles River (Wilmington, Massachusetts). They were acclimatized in a controlled environment at the Rutgers University Newark Animal Facility for at least 7 days, with a 12-hour light-dark cycle, 22 ± 1 °C temperature, and ad libitum access to food and water. All experiments were performed with Rutgers Institutional Animal Care and Use Committee (IACUC) approval (PROTO202100043).

Dams at 8 weeks of age were bred 2:1 and plugs were checked twice a day. Gestational day (GD) 0.5 is the day at which plugs were detected and GD 1 was the following day. Mice were orally dosed via gentle pipet feeding starting at GD 8 until GD 15 in three treatment groups. The treatment groups were vehicle control (ultrapure water), 5 mg/kg of 50 nm blue fluorescent nanoplastics, or 5 mg/kg of 200 nm red fluorescent polystyrene nanoplastics. Mice were euthanized on GD 15 by CO_2_ following the IACUC-approved protocol.

Placenta and fetus weights were recorded after euthanization. Placentas were cut in half, with one half snap frozen in liquid nitrogen for histomorphology and the other half fixed in 4% paraformaldehyde overnight and transferred to 70% ethanol for immunohistochemistry staining and imaging.

The sizes of the particles (50 and 200 nm) were chosen to be small enough to be expected to translocate to the placenta and large enough to be expected to be detectable using fluorescent microscopy [9,10]. The dose of 5 mg/kg was chosen to fall within an estimated environmentally relevant range and to compare to previous studies. At the time this experiment was designed, we used the estimate of human ingestion of 0.1–5 g of microplastics per week, which would correspond to 0.23 – 11.9 mg/kg/day of plastic intake for an adult woman with an average mass of 60 kg [29]. This assessment has been criticized as an overestimation [30,31]; however, body surface area conversions for mice, assuming a 20 g mouse, suggest that the mouse equivalent exposure would be 2.5 – 145 mg/kg [21,32–35].

### 2.3 Fluorescent microscopy

High-resolution fluorescence images of nanoparticle distribution within the placenta were acquired using a Keyence microscope (BZ-X800) and image stitching function of the BZ-X analyzer.

### 2.4 Morphological analysis

Half placentas from CD-1 mice were fixed in 4% paraformaldehyde overnight and transferred to 70% ethanol. Placentas were embedded in paraffin wax and sectioned at 5 μm. Sections were mounted on glass slides and stained with hematoxylin and eosin (H&E). Coverslipped slides were digitized using a Zeiss Axioscan7. Randomly selected placentas from at least one male and one female fetus were used per dam for analysis.

Placentas were analyzed for two histological outcomes using the Zeiss Zen software: placental zone area and maternal and fetal blood spaces. One central section from each placenta was used for all measurements. The placentas were first measured for changes in area of three key placental zones: the labyrinth zone, the basal zone, and the decidua. The layers were identified by comparing our samples to placenta morphology in previous publications [36]. We traced the outline of each layer using the contour spline tool. Layers were analyzed for changes in the total area by dam and by sex.

We also measured the areas of maternal and fetal blood spaces (MBS and FBS, respectively) using the contour spline tool and the length of the intrahemal barrier (IHB) using the line feature. Maternal blood spaces can be identified by small red blood cells that clump together, and they are generally much larger. Fetal blood spaces are much smaller and contain 1-2 much larger cells. Fetal red blood cells may or may not have shed their nucleus. For each section, we measured the MBS, FBS, and IHB three times and averaged the measurements. Blood spaces were analyzed for changes in area and length by dam and by sex.

### 2.5 Gene expression

Total RNA was extracted from at least one randomly selected frozen placenta half per sex per dam using E.Z.N.A. MicroElute® Total RNA Kit according to the manufacturer’s protocol. The RNA concentration was quantified using Thermo Scientific Nanodrop One C Spectrophotometer. For cDNA synthesis, iScript Reverse Transcription kit was used. Quantitative PCR was conducted using 94°C for 30s, 42 cycles at 94°C for 5s, and 60°C for 34s cycling conditions. The genes that were analyzed were (**Table 1**) smad family number 2 (*Smad2*), smad family number 3 *(Smad3*), transforming growth factor beta 1 (*Tgfb1*), activin a receptor type 2a (*Acvr2a*), estrogen receptor 1 (*Esr1*), cadherin 1 (*Cdh1*), cadherin 11 (*Cdh11*), cholesterol side-chain cleavage enzyme (*Cyp11a1*), prostaglandin-endoperoxide synthase 2 (*Ptgs2*), prolactin related genes (*Prl7d1*, *Prl3d1,* and *Prl3b1*), angiotensin I converting enzyme (*Ace*), angiotensin converting enzyme 2 (*Ace2*), matrix metallopeptidase 10 (*Mmp10*), fibroblast growth factor binding protein 1 (*Fgfbp1*), cofilin 1 (*Cfl1*), epidermal growth factor receptor (*Egfr*), placental growth factor (*Pgf*), growth differentiation factor 15 (*Gdf15*), Sry-box transcription factor 9 (*Sox9*), and peroxisome proliferator activated receptor gamma (*Pparg*). qPCR analysis was done using the Pfaffl method with the housekeeping gene beta-actin (*ActB*) as the reference gene [37]. Specificity of primers was confirmed using melting curve analysis and genes with primers that had multiple significant melting curve peaks were excluded from the study. Sex of placentas for all analyses was determined by cross-referencing the genes for sex determining region of Chr Y (*Sry*) and inactive X specific transcripts (*Xist*).

**Table 1:**
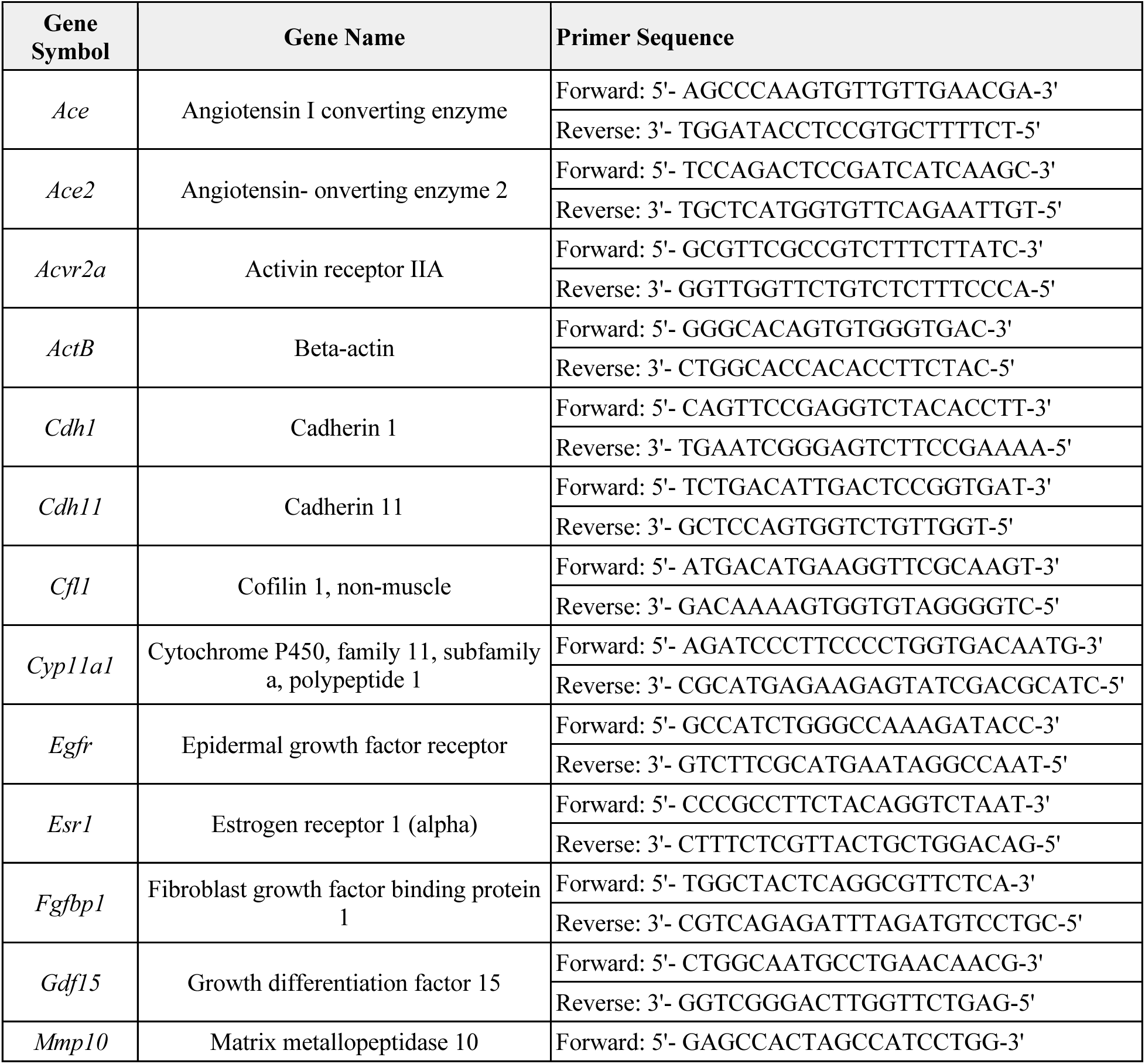
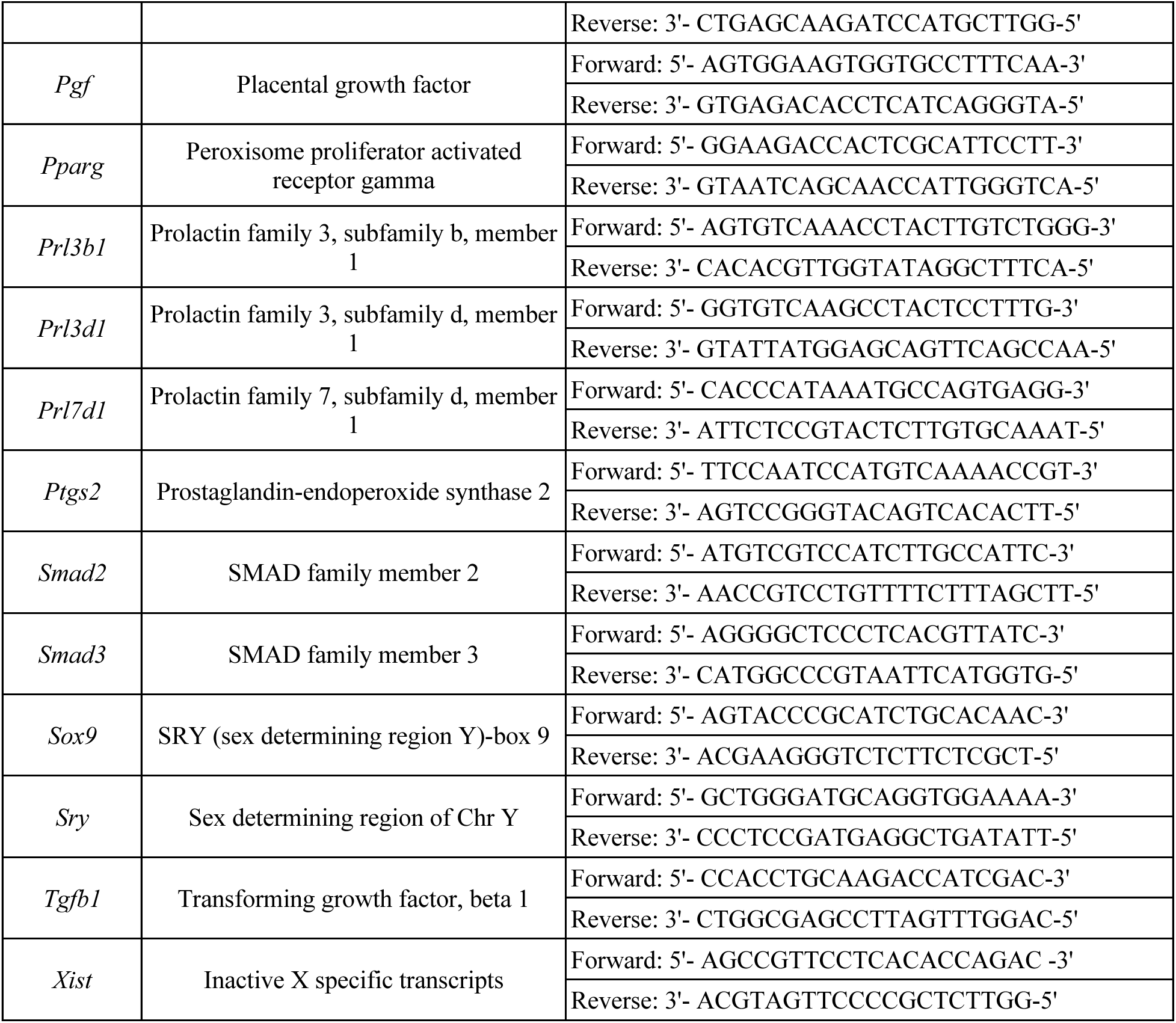
Primer sequences.

### 2.6 Statistical analysis

Data are expressed as means ± standard error of the mean (SEM) by dam for 3-5 dams per treatment group. Data were analyzed by comparing treatment group to control using IBM SPSS version 29 software (SPSS Inc., Chicago, IL, USA). All data were continuous and assessed for normal distribution by Shapiro-Wilk analysis. If data met assumptions of normal distribution and homogeneity of variance, data were analyzed by one-way analysis of variance (ANOVA) followed by Tukey HSD or Dunnett 2-sided post-hoc comparisons. However, if data met assumptions of normal distributions, but not homogeneity of variance, data were analyzed by ANOVA followed by Games-Howell or Dunnett’s T3 post-hoc comparisons. If data were not normally distributed or presented as percentages, the independent sample Kruskal-Wallis H followed by Mann-Whitney U non-parametric tests were performed. For all comparisons, statistical significance was determined by p-value ≤ 0.05. If p-values were greater than 0.05, but less than ≤ 0.1, data were considered to exhibit a trend towards significance.

## 3 Results

### 3.1 Particle characterization

Nanoplastic particles were characterized to confirm shape and size. Particles purchased as 200 nm were determined to have an actual average size of 220 nm with a polydispersity index (PDI) of 0.085 [38]. Purchased 50 nm particles had an average diameter of 48.2 nm with a PDI of 0.049 (**Figure 1**).

**Figure 1:**
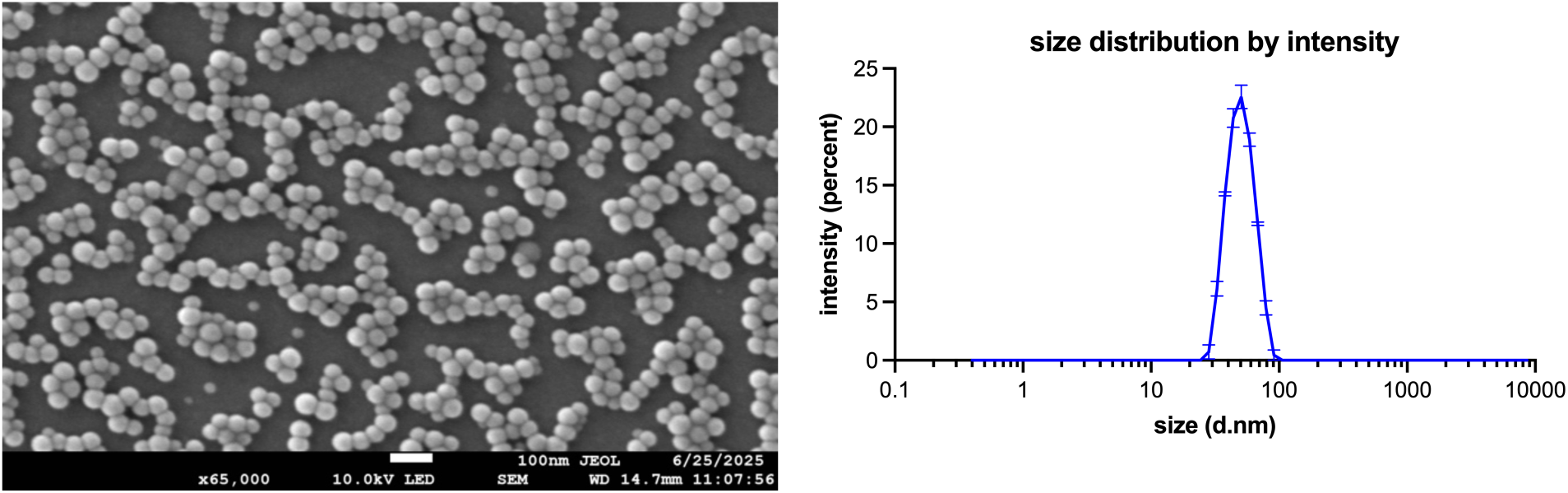
50 nm polystyrene spheres used in this study were characterized by scanning electron microscopy (SEM, scale bar 100 nm, left) and dynamic light scattering (DLS, right). Characterization of 200 nm polystyrene spheres was previously reported [38].

### 3.2 Confirmation of plastic distribution to the placenta

Fluorescent imaging of placenta sections confirms that maternal oral exposure to nanoplastics results in plastic translocation to the placenta (**Figure 2).**

**Figure 2:**
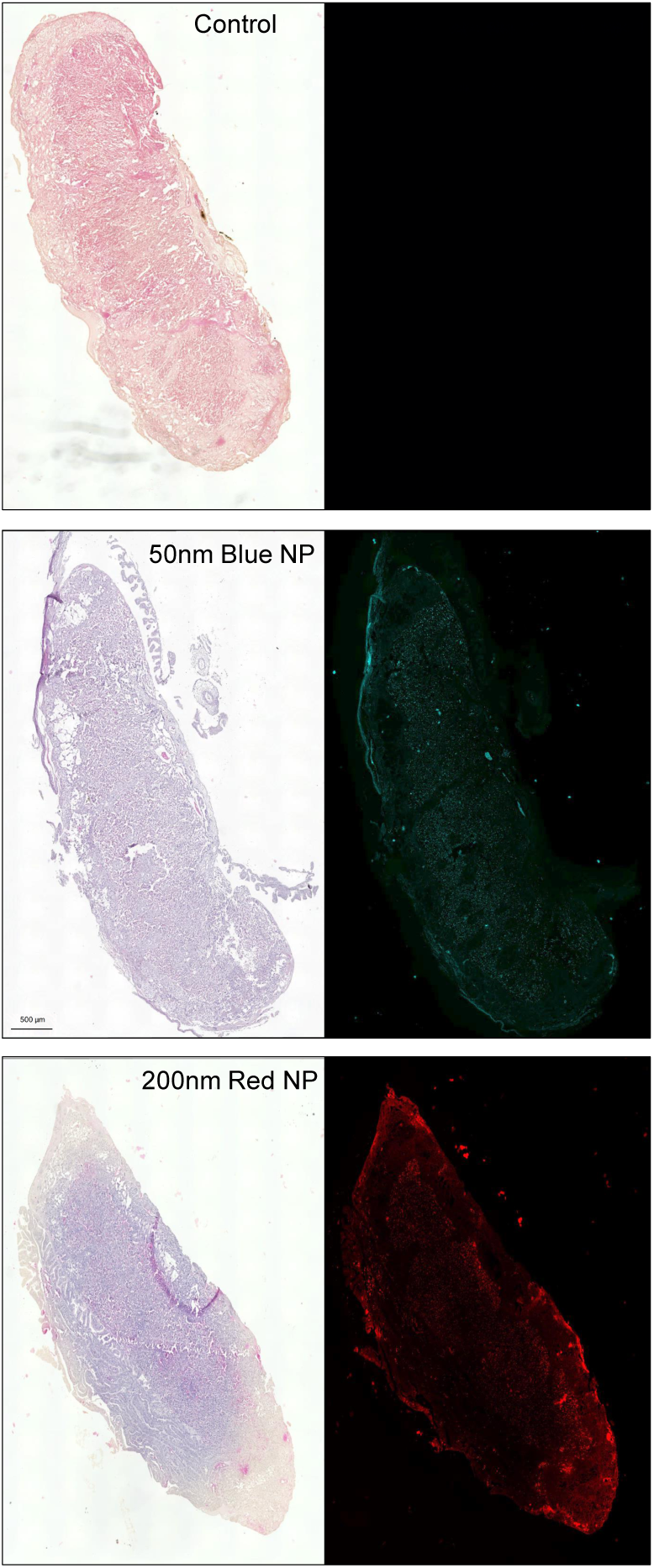
Hematoxylin and eosin-stained slides from each treatment group were imaged to confirm that the fluorescent particles are present in placentas from exposed animals and not controls.

### 3.3 Organ Weights

Placental and fetal weights were recorded and used to calculate the placenta:fetus weight ratio, which are reported per dam (**Figure 3**). No significant differences were observed in treatment groups compared to control when sexes were combined or separated

**Figure 3:**
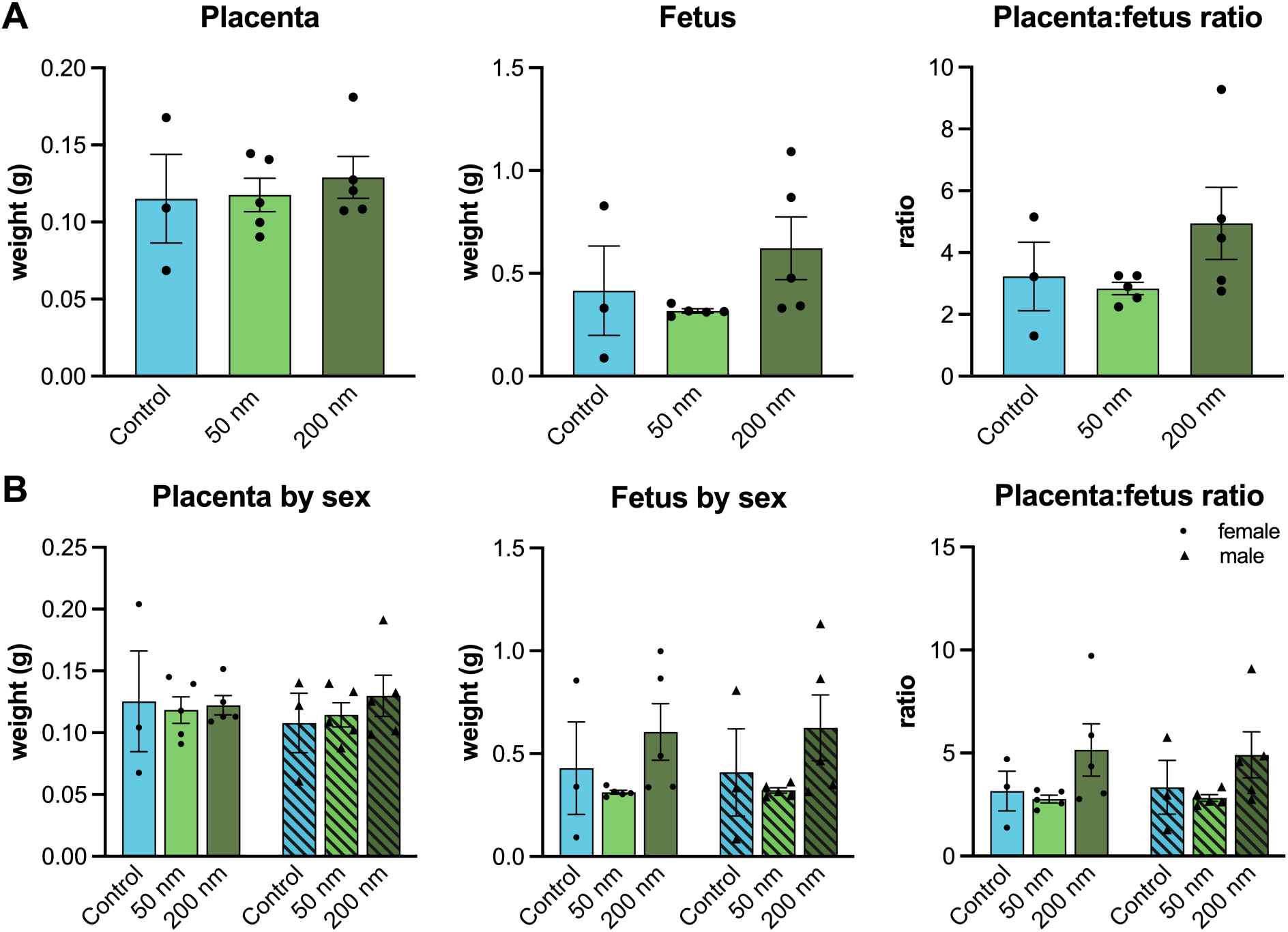
Impacts of nanoplastic exposure on placenta weight, fetus weight, and placenta:fetus weight ratio after oral exposure to 5 mg/kg PS NP from GD 8-15. Results are reported per dam (A) and for each sex per dam (B).

### 3.4 Morphology analysis

The placentas were measured for changes in area of the labyrinth zone, the basal zone, and the decidua. The labyrinth is the innermost layer on the fetal side and is highly vascularized with branched villi to transport nutrients and wastes to and from the fetus. The labyrinth has clearly identifiable and measurable maternal and fetal blood spaces. The middle layer is the basal zone, also known as the junctional zone, which forms the invasive interface between the labyrinth and maternal decidua. The basal zone contains invasive fetal trophoblast cells, including spongiotrophoblasts and glycogen-containing trophoblasts. Finally, the decidua is maternal derived, containing uterine decidual cells and maternal blood vessels (**Figure 4A**).

**Figure 4:**
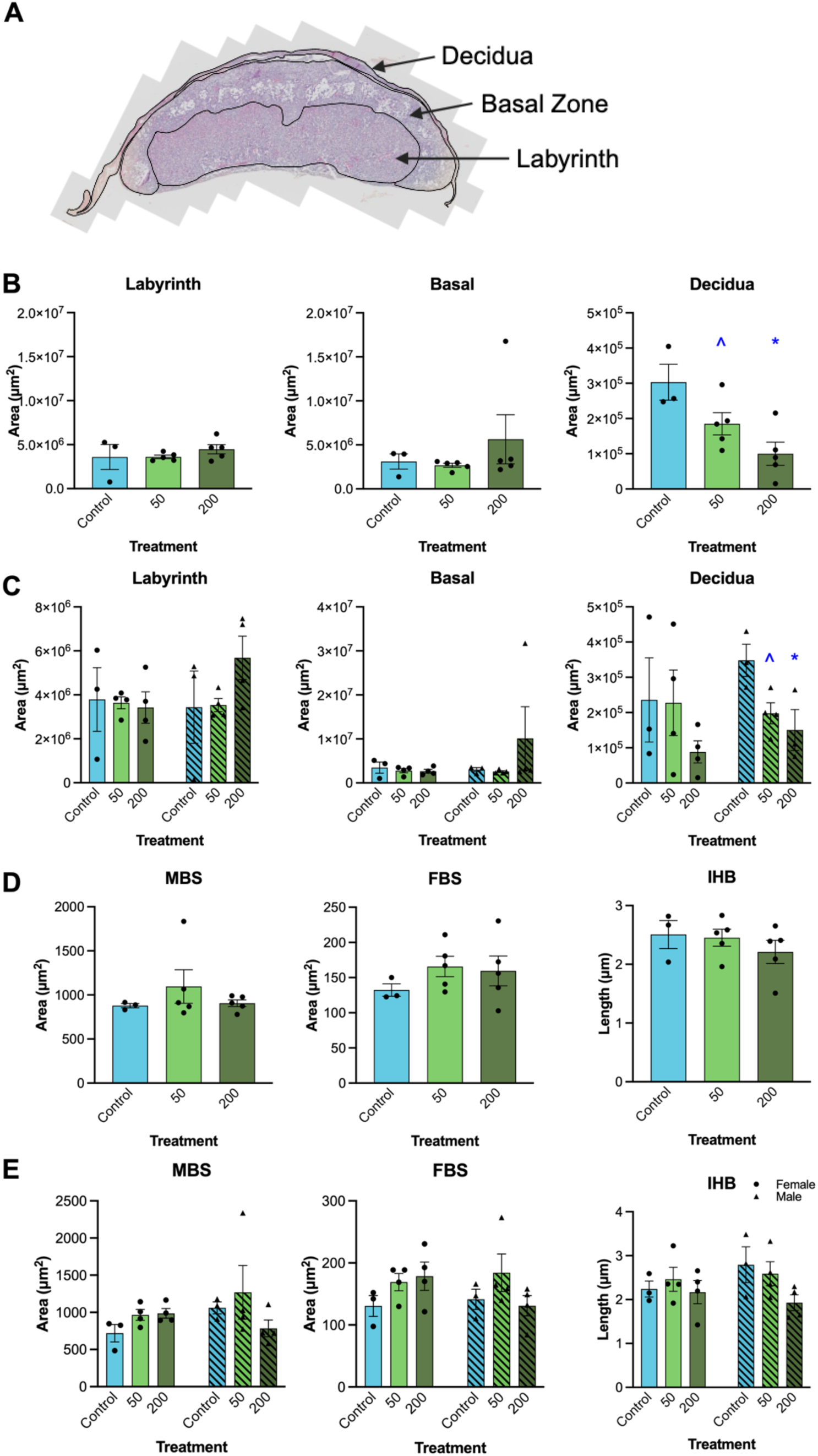
Impact of nanoplastics on placenta morphology. Mid placenta sections (A) from at least two placentas per dam exposed to 50 nm or 200 nm polystyrene nanoplastics from gestational day 8-15 were stained with hematoxylin and eosin and measured for placenta layer area by dam (B) and by sex (C). Area of maternal blood spaces (MBS) and fetal blood spaces (FBS) and length of intrahemal barrier (IHB) were measured by dam (D) and by sex (E). Graphs represent mean ± SEM from 3-5 dams per treatment group. Asterisks (∗) indicate significant differences from the control (p ≤ 0.05) and ^ indicates a trend toward significance (p ≤ 0.10).

The area of the decidua was significantly decreased in the 200 nm treatment group compared to control (p = 0.008) and borderline decreased in the 50 nm treatment group compared to control (p = 0.097) (**Figure 4B**). When layers were analyzed by sex, only male placentas were impacted. Decidua area was significantly decreased in the 200 nm treatment group compared to control (p = 0.048) and borderline decreased in the 50 nm treatment group compared to control (p = 0.075) (**Figure 4C**). Maternal blood space area, fetal blood space area, and intrahemal barrier were not significantly different compared to control placentas either overall or by sex (**Figure 4D-E**).

### 3.5 Gene Expression

We performed qPCR on a panel of genes related to TGFB signaling and placenta function, including direct members of the TGFβ pathway (*Smad2, Smad3, Tgfb1, Gdf15, Acvr2a*), steroid hormones and metabolic regulators (*Esr1, Pparg, Cyp11a1*), lactogens (*Prl3d1, Prl7d1, Prl3b1*), adhesion and remodeling (*Cdh1, Cdh11, Cfl1, Mmp10, Sox9*), and vascularization and angiogenesis (*Egfr, Fgfbp1, Pgf, Ace, Ace2, Ptgs2*). First, we constructed a heatmap to show all genes investigated to looks for trends by class of gene and exposure. At first glance, more genes appear to be downregulated in the 50 nm treatment group compared to the 200 nm treatment group in all samples (**Figure 5A**). When samples were split by sex, more and greater magnitude of gene expression changes are evident in males placentas compared to female placentas (**Figure 5B**).

**Figure 5:**
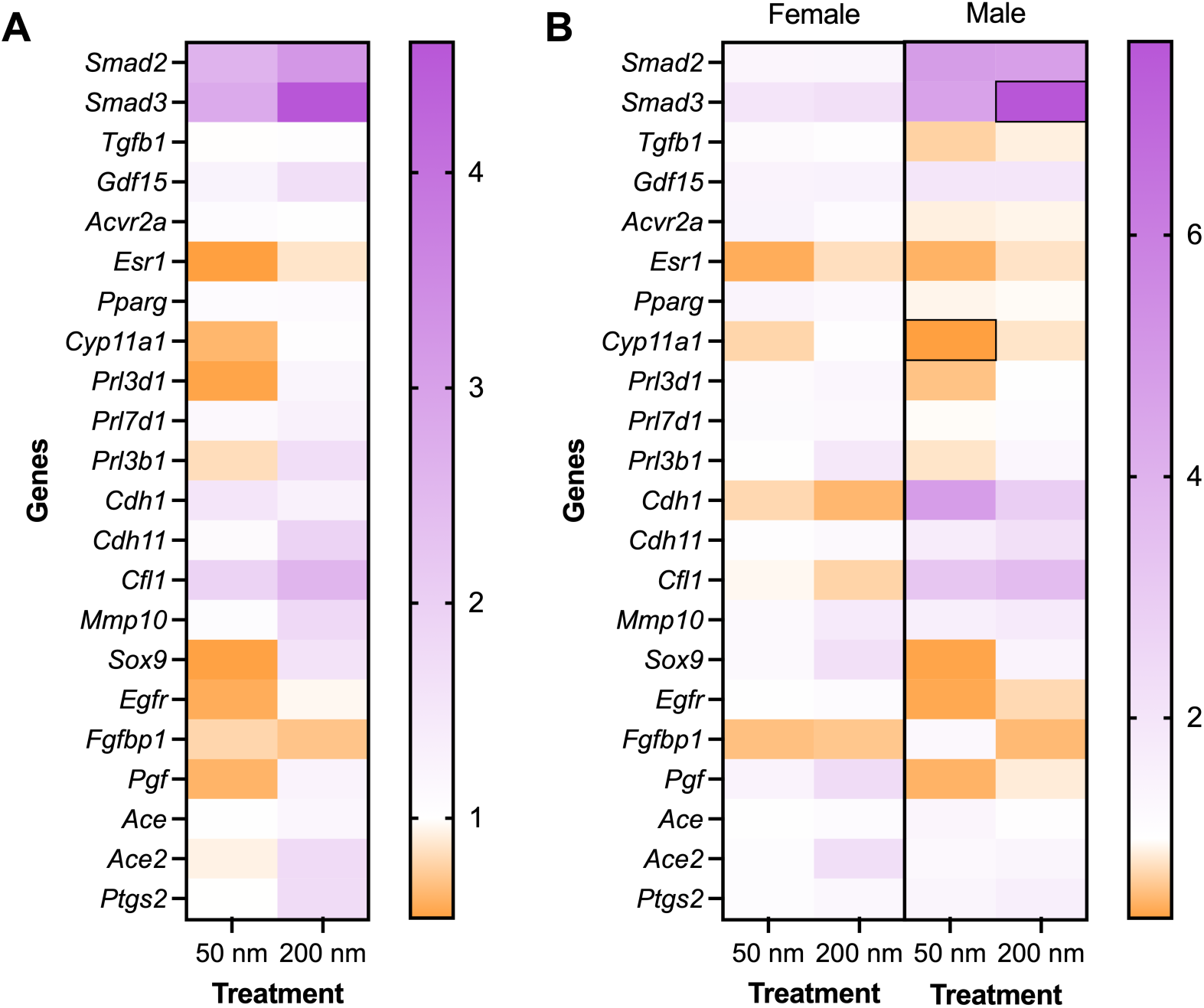
Heatmap of relative gene expression compared to control in mouse placentas exposed to 50 nm or 200 nm polystyrene nanoplastics by dam (A) and by sex (B). Black boxes indicate statistically significant results at p < 0.05. Graphs below show individual data points and note trending (p ≤ 0.10) results, as the sample size was low, limiting power.

#### 3.5.1 TGFβ pathway

In the TGFβ signaling pathway, *Tgfb1* codes for a ligand for the TGFβ family of receptors, which recruit and activate SMAD family transcription factors with widespread downstream targets. GDF15 is a TGFβ family protein. GDF15 promotes invasion of trophoblast cells, and decreased levels of GDF15 in human serum is associated with miscarriage. The activin receptor ACVR2A is another example of a TGFβ superfamily protein. In placentas from mice exposed to 50 or 200 nm polystyrene nanoplastics for 7 days during placenta formation, transcripts for *Smad3* in the 50 nm treatment group (p = 0.057) and *Smad2, Smad3*, and *Gdf15* were increased in the 200 nm treatment group (p = 0.069, 0.075, and 0.101, respectively) compared to control (**Figure 6A**). When analyses were separated by placenta sex, *Smad3* was significantly upregulated in males in the 200 nm treatment group (p = 0.045) and *Tgfb1* trended towards downregulation in the 50 nm treatment group (p = 0.107) compared to control (**Figure 6B**).

**Figure 6:**
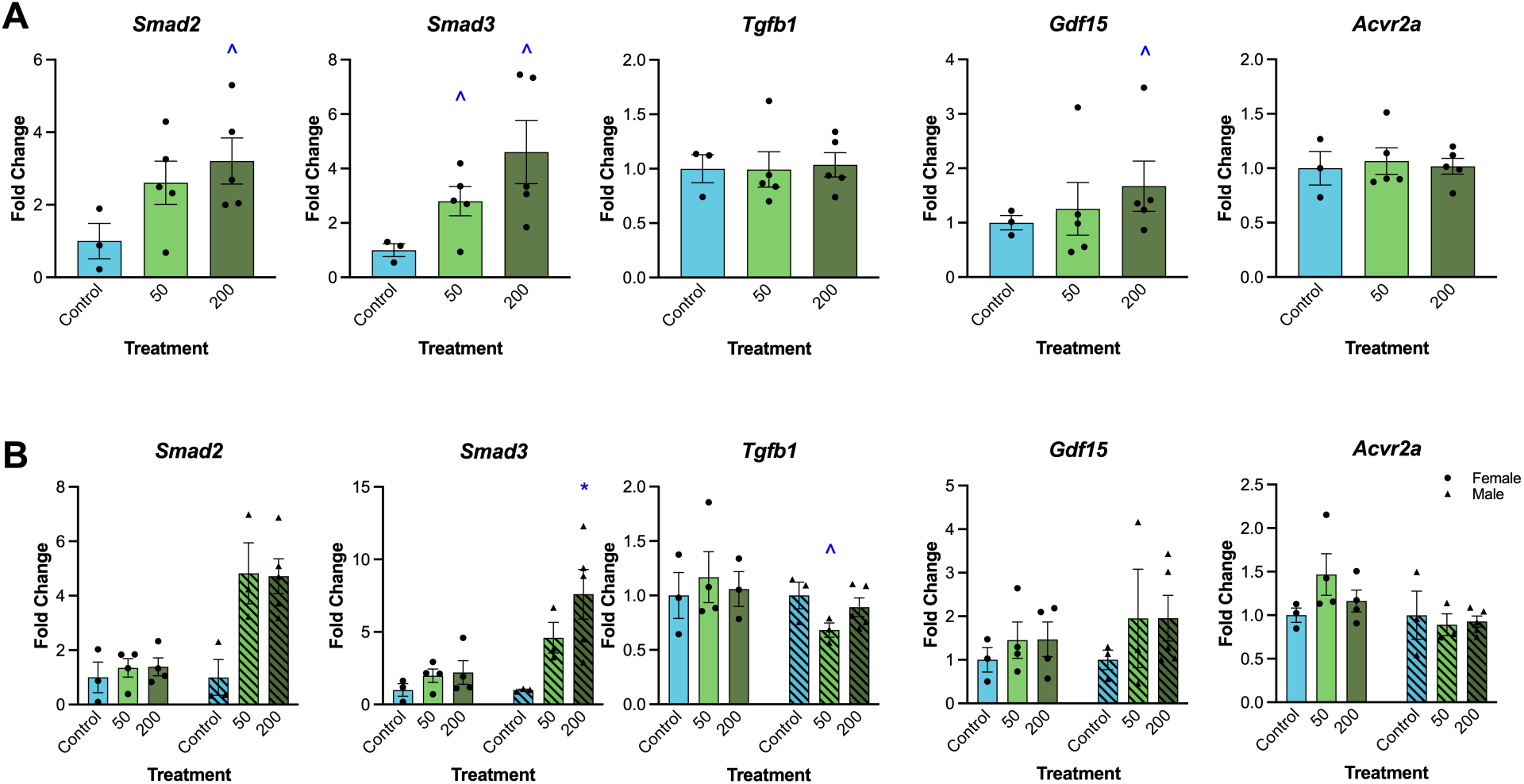
Impact of nanoplastic exposure on placenta expression of genes related to the TGFβ pathway by dam (A) and by sex (B). Graphs represent mean ± SEM from 3-5 dams per treatment group. Asterisks (∗) indicate significant differences from the control (p ≤ 0.05) and ^ indicates a trend toward significance (p ≤ 0.107).

#### 3.5.2 Steroid Hormones

We measured expression of genes for estrogen receptor alpha (*Esr1*) and peroxisome proliferator-activated receptor gamma (*Pparg*), nuclear receptors with important functions in the developing placenta, and cytochrome P450 family 11 subfamily A member 1 (*Cyp11a1*), which encodes for the enzyme that converts cholesterol to pregnenolone in the steroidogenesis pathway. We were unable to measure *Esr2*, *Ar*, or *Cyp19a1* because expression levels were too low for quantitation. Expression of *Esr1* was borderline decreased (p = 0.053) in the 50 nm treatment group compared to control in the overall samples (**Figure 7A**) and in males only (p = 0.097, **Figure 7B**). *Cyp11a1* was significantly decreased in males only at 50 nm (p = 0.005), whereas *Pparg* was borderline significantly increased in females only in the 50 nm treatment group (p = 0.083) compared to control.

**Figure 7:**
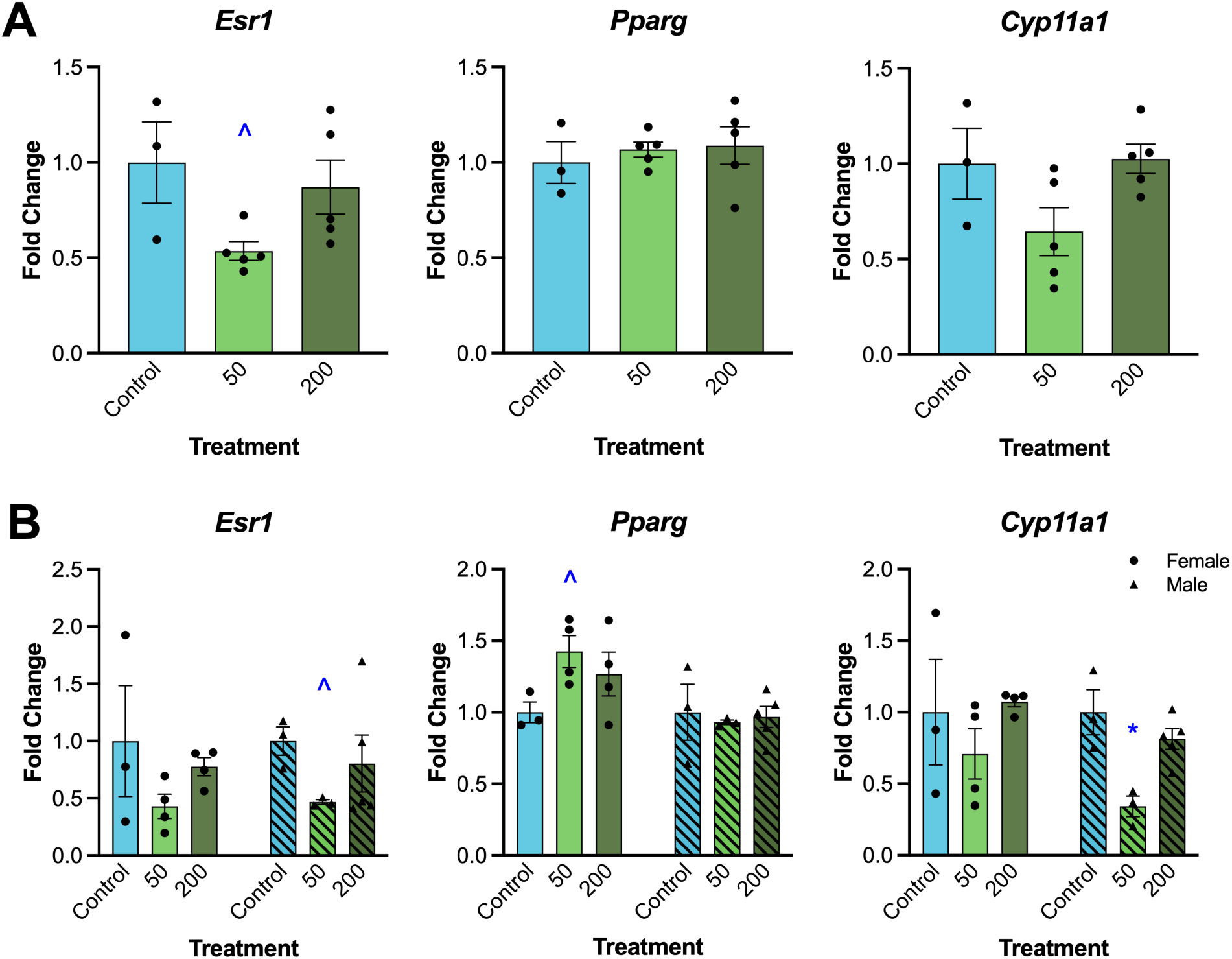
Impact of nanoplastic exposure on placenta gene expression related to steroid hormones by dam (A) and by sex (B). Gene expression was measured via qPCR. Graphs represent mean ± SEM from 3-5 dams per treatment group. Asterisks (∗) indicate significant differences from the control (p ≤ 0.05) and ^ indicates a trend toward significance (p ≤ 0.10).

#### 3.5.3 Lactogens

We measured expression of three placenta lactogens, *Prl3b1, Prl3d1,* and *Prl7d1*, members of the prolactin hormone family. Expression of *Prl3d1* was borderline significantly decreased (p = 0.10) in the 50 nm treatment group compared to control, but no trends were observed when samples were separated by sex (**Figure 8**).

**Figure 8:**
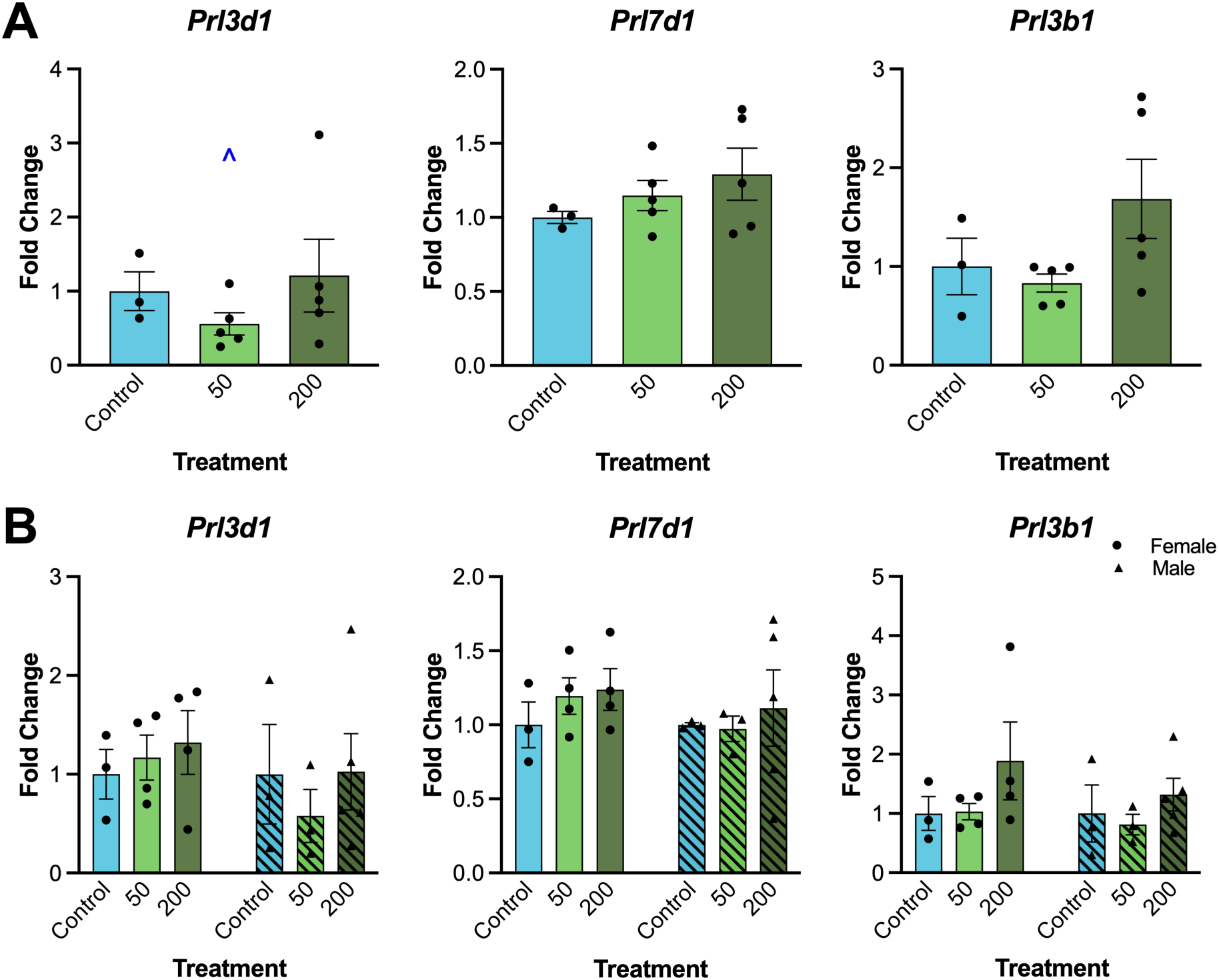
Impact of nanoplastic exposure on placenta gene expression related to lactogens by dam (A) and by sex (B). Gene expression was measured via qPCR. Graphs represent mean ± SEM from 3-5 dams per treatment group. Asterisks (∗) indicate significant differences from the control (p ≤ 0.05) and ^ indicates a trend toward significance (p ≤ 0.10)

#### 3.5.4 Adhesion, Remodeling, Growth, Angiogenesis, and Vascularization

From a panel of genes related to the essential placenta functions of adhesion, remodeling, growth, angiogenesis, and vascularization in the developing placenta, *Cofilin 1* (*Cfl1)* was borderline significantly upregulated in the 50 nm treatment group compared to control (p = 0.10), but no trends were observed when samples were separated by sex (**Figure 9**). Conversely, placenta growth factor (*Pgf*) was borderline significantly increased in females in the 200 nm treatment group only (p = 0.056) compared to control (**Figure 10**).

**Figure 9:**
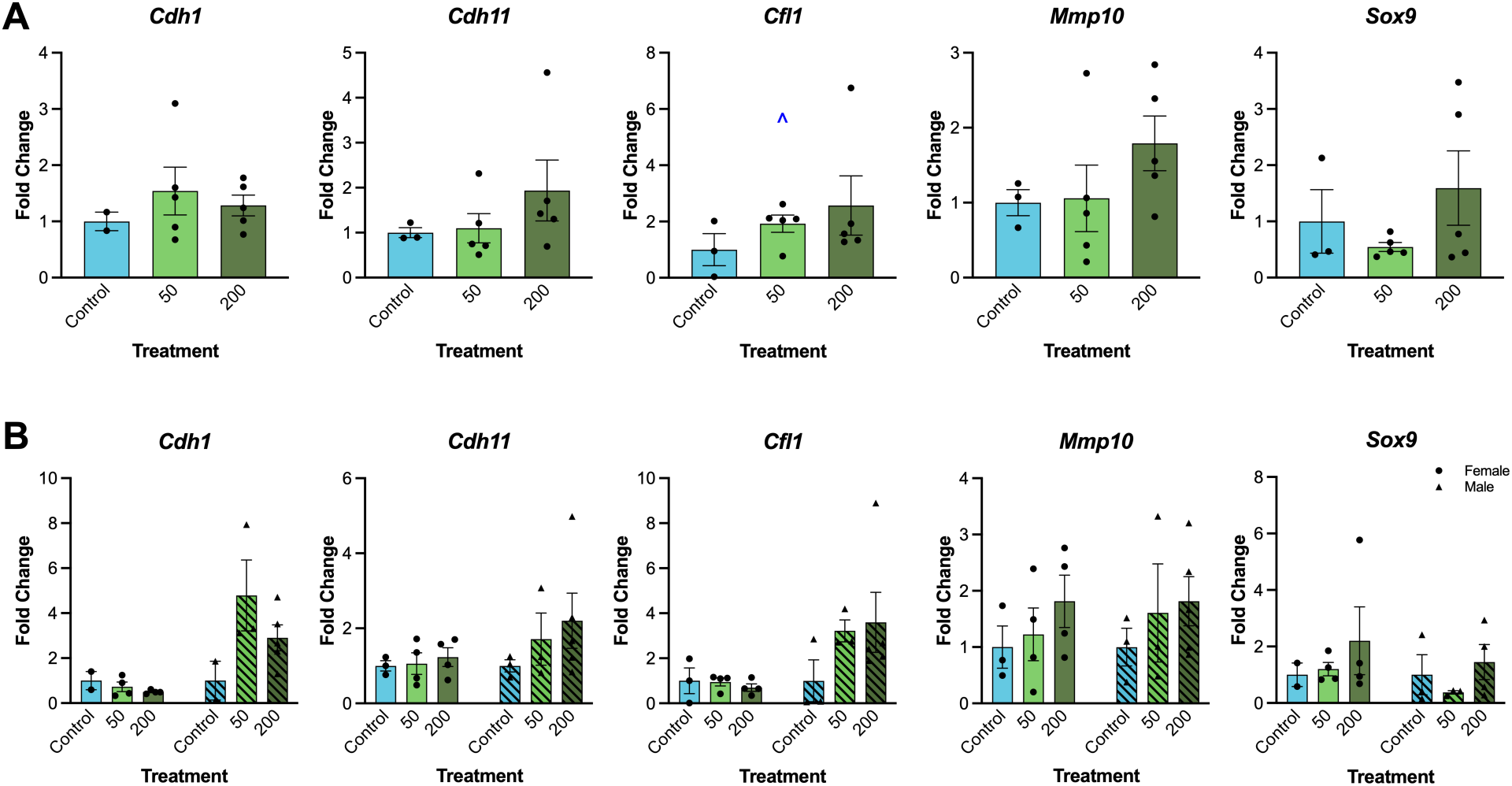
Impact of nanoplastic exposure on placenta gene expression related to adhesion and remodeling by dam (A) and by sex (B). Gene expression was measured via qPCR. Graphs represent mean ± SEM from 3-5 dams per treatment group. Asterisks (∗) indicate significant differences from the control (p ≤ 0.05) and ^ indicates a trend toward significance (p ≤ 0.10).

**Figure 10:**
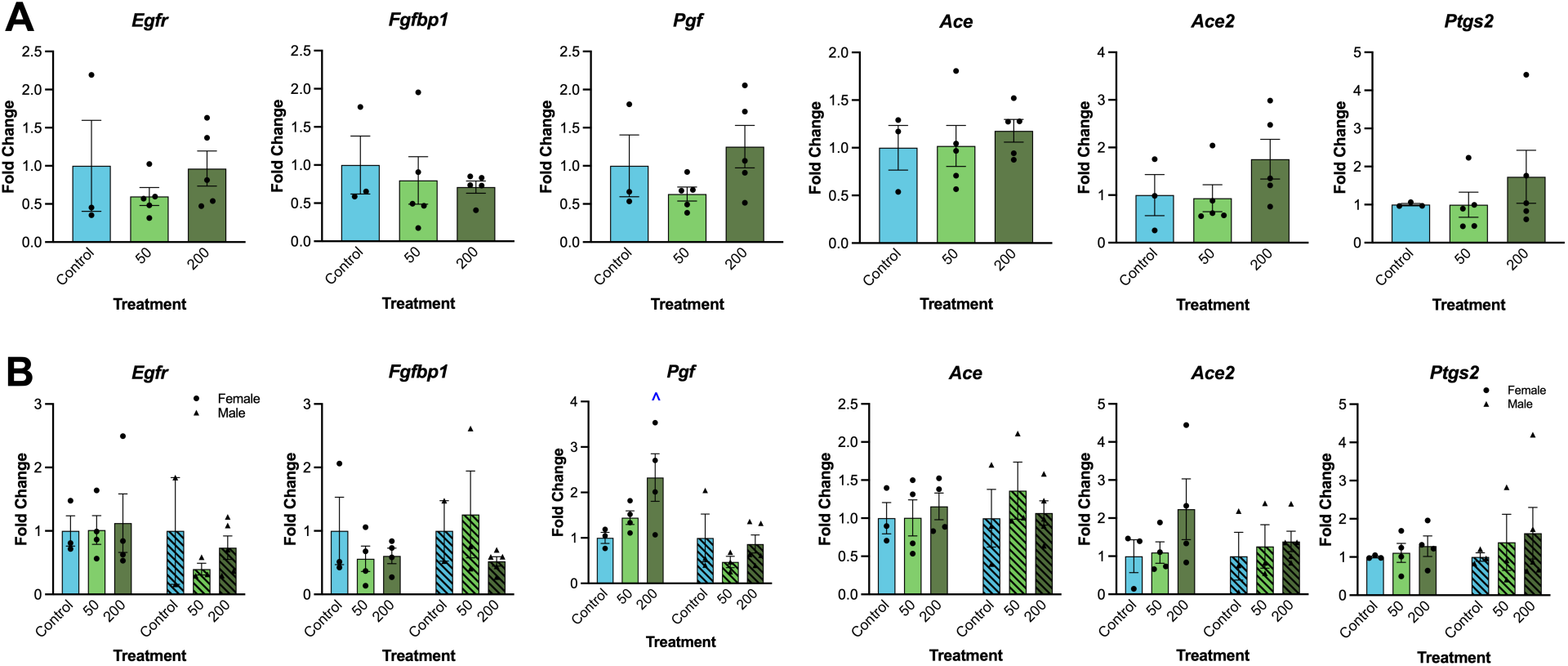
Impact of nanoplastic exposure on placenta gene expression related to growth, angiogenesis, and vascularization by dam (A) and by sex (B). Gene expression was measured via qPCR. Graphs represent mean ± SEM from 3-5 dams per treatment group. Asterisks (∗) indicate significant differences from the control (p ≤ 0.05) and ^ indicates a trend toward significance (p ≤ 0.10).

## 4 Discussion

Disruption of placenta development or function at the molecular level can result in significant pregnancy complications such as placenta accreta, preeclampsia, and restricted placenta size [17]. Multiple studies have characterized MNPs in human placentas and other tissues, including meconium, which indicates fetal exposure [5,6,39,40]. Microplastics in human placentas have been associated with low birth weight in pregnancies characterized by IUGR, a pregnancy complication related to placental insufficiency and affecting 10-15% of pregnancies worldwide [14]. A large prospective cohort recently found higher placenta microplastics were associated with reduced birth weight, birth length, and head circumference [15]. Previous rodent studies suggest that NPs may contribute to pregnancy complications related to reduced placenta size [41]. In this work, we tested the hypothesis that nanoplastic exposure during the period of placenta formation would disrupt placenta morphology and signaling, specifically focusing on TGFβ signaling, an understudied potential mechanism of action. Overall, we identified disruption to placenta layer area and signaling pathways involved in placenta development.

We previously confirmed presence of the red 200 nm particles in placentas from this cohort of exposed animals using enhanced darkfield hyperspectral microscopy (CytoViva) and flame ionization mass spectrometry [42]. In this study, we visually confirmed the presence of both sizes of particles using fluorescent microscopy. The translocation of particles into the placenta from prenatal exposure is consistent with previous reports from late gestational inhalation exposure in rats [7], oral gavage in rats [43], late gestation intravenous injection in mice [9], and oral gavage in mice [13,35,44].

The overall size (weight or area) of the placenta can indicate placenta insufficiency that may decrease its ability to support the fetus, leading to intrauterine growth restricted fetuses or pregnancy complications. In one of the few human epidemiology studies focusing on MNPs, the presence of particles in human placentas was associated with decreased fetal size in IUGR pregnancies [14], emphasizing the need for further study. Examining placenta layer area in controlled animal studies can provide insight into MNP targets, as each of the three layers of the placenta play important roles and have visually distinguishable phenotypes [45–47]. In mice, the placenta is fully developed around day 12.5 and reaches peak weight around day 15.5 [47]. Around this time, the decidua begins to thin in preparation for birth. In this study, we did not find an impact in overall placenta weight or efficiency or the sizes of maternal and fetal blood spaces, but we did identify a decrease in the area of the decidua in both nanoplastics treatment groups, which appears to be mostly male driven. From the single timepoint of collection, it is not clear if small decidua were underdeveloped, which can lead to placenta insufficiency and IUGR, or developed normally but prematurely began the thinning process. Under-decidualization and thin decidua are hallmarks of aging [48]. Phthalate exposure earlier in gestation has been shown to impact decidualization and increase the proportion of junctional zone area compared to labyrinth area at gestational day 13, but the area of the decidua was not assessed [49].

A few previous studies have identified disruption of placenta morphology from MNP exposure, although no others have identified direct changes in the decidua, which may be due to differences in particle properties, exposure window, or collection timepoint. Mice exposed to 0.1 mg/day (∼5 mg/kg) of 50-100 nm PSNPs via drinking water throughout gestation had decreased total placenta area and labyrinth area at gestational day 12.5 and 18.5, although this study did not characterize their particles nor analyze changes in decidua area [21]. Pregnant mice exposed by gavage every three days to nanoplastics extracted from plastic lined beverage cups using hot water showed a dose-dependent decrease in labyrinth area on day 18.5 [35]; other layer areas were not reported. Rats exposed orally via pipet to 1 mg/kg of 100 nm PSNPs from gestational day 5-20 had decreased placenta weight and increased placenta efficiency in males only compared to control, but no significant differences in the decidua, basal, or labyrinth layer area [19].

We measured expression of a panel of *a priori* selected genes related to TGFB signaling and placenta function, including regulators of the TGFβ pathway, nuclear receptors and sex steroid hormones, lactogens, and adhesion, remodeling, vascularization, and angiogenesis factors largely under TGFβ transcriptional control. Power for this pilot project was limited due to small sample size, so we considered overall trends as well as statistically significant effects. Gene expression results overall show a mix of up and down regulated genes, with a trend toward more genes downregulated in the 50 nm treatment group and more genes upregulated in the 200 nm treatment group. By sex, male placentas had more and greater magnitude gene expression changes compared to female placentas, although individual statistically significant results do not show a clear pattern.

During pregnancy, the TGFβ superfamily is critical for implantation, decidualization, maternal-fetal tolerance, and overall placental development in humans [50]. In canonical TGFβ signaling, hormones such as transforming growth factor betas and activins serve as ligands for receptors to activate mediators in the SMAD family via phosphorylation, which then serve as transcription factors [24]. The ligand *Tgfb1* and the activin receptor *Acvr2a* were not significantly disrupted, but *Gdf15* trended toward increased expression. *Smad2* and *Smad3*, which are under TGFB and activin control, were increased from exposure to both NP sizes. Increased expression of *Smad2/3* may be a compensatory response for disrupted phosphorylation and furthermore suggests that downstream transcriptional control may be impacted. We observed a trending increase in *Cfl1*, which is thought to be under Smad-independent TGFβ control, in the 50 nm treatment group and a trending increase in *Pgf* in females in the 200 nm treatment group, but other TGFβ transcriptionally controlled genes in Figure 9 and 10 were not impacted, and no overall trend was observed. However, the decrease in expression of the prolactin gene *Prl3d1* may be related to increased SMADs, as previous studies indicate that prolactins are inhibited by TGFβ signaling, specifically through SMAD3 [51]. In addition, SMAD3 is highly localized within the decidua during decidualization [52]. The decreased decidua area in the present study in exposed animals compared to controls may indicate defective decidualization, mediated by TGFβ/SMAD dependent signaling, ultimately leading to a reduction in total decidual stromal fibroblasts.

Similarly to prolactins, estrogen and TGFβ signaling have opposing interactions. TGFβ1 inhibits aromatase in JEG3 cells via Smad2 phosphorylation [53], and a recent report in human bronchial epithelial cells showed that TGFB1 suppressed both gene and protein expression of ESR1 [54]. Similarly, we observed a decrease in *Esr1* in the 50 nm treatment group, which may be either the cause of or a compensatory effect related to increased *Smad2/3*. As other studies suggest that MNP exposure may alter steroid hormone production [38,55], including in the placenta [21], we also measured *Cyp11a1* expression, which converts cholesterol to pregnenolone. Our observed decrease in *Cyp11a1* expression is consistent with another recent report in which mice exposed to PS MNPs in a range of sizes had significantly decreased levels of *Cyp11a1* at both GD12.5 and 18.5 [21], providing further evidence that NPs are potential endocrine disruptors.

This study is the first to explore the potential for nanoplastic exposure to disrupt TGFβ signaling in the placenta, a key modulator of many placenta functions. The timing of exposure was selected to assess the period of placenta formation without interfering with implantation. As an exploratory pilot study, weaknesses include small sample size that limited power to detect all but large differences between treatment groups and limited selection of targets in the pathways of interest. Future studies will use more environmentally relevant nanoplastic types, including other polymers and particle shapes. We are currently working to quantify concentrations of each plastic in the placental tissues [42].

## 5 Conclusion

In conclusion, gestational exposure to polystyrene nanoplastics disrupted placenta morphology and altered gene expression of signaling modulators in the TGFβ pathway at gestational day 15 in mice. In addition, nuclear receptor and steroidogenic enzyme gene expression was altered. We observed sex specific differences, with morphology more disrupted in male placentas than female. Effects were not consistent across nanoplastics sizes. Overall, these results support the need for further study of nanoplastic interactions with TGFβ signaling in the placenta and other tissues and contribute to the growing body of evidence suggesting that nanoplastics can act as endocrine disruptors.

## 8 Declaration of generative AI and AI-assisted technologies in the manuscript preparation process

Generative AI was not used in the preparation of this article.

## 9 Author CRediT Statement

Hanin Alahmadi: conceptualization, investigation, supervision, project administration, writing – original draft; Allison Harbolic: investigation, methodology, visualization, validation; Christopher De Oliveira-Cordova: investigation, supervision, writing – original draft; Raulle Reynolds: data curation, validation, visualization; Michelle Jojy: investigation, visualization; Courtney Potts: investigation; Sofia Doan: investigation, visualization; Tanvi Mathur investigation, visualization; Mohammad Saiful Islam: investigation, visualization; Melisa Andrade: investigation; Quinton Smith: methodology, supervision; Phoebe A. Stapleton: methodology, supervision; Somenath Mitra: funding acquisition, resources, supervision; Genoa R. Warner: conceptualization, data curation, formal analysis, funding acquisition, investigation, methodology, project administration, supervision, validation, writing – original draft, writing – review & editing

## 10 Acknowledgements

Thank you to members of the EDC Lab for help with animal experiments. AH thanks Chelsea Cary and Destiny McWilliams for methods training.

## References

[1] R. Geyer, J.R. Jambeck, K.L. Law, Production, use, and fate of all plastics ever made, Sci Adv 3 (2017) e1700782. 10.1126/sciadv.1700782.

[2] J. Gigault, A.T. Halle, M. Baudrimont, P.-Y. Pascal, F. Gauffre, T.-L. Phi, H. El Hadri, B. Grassl, S. Reynaud, Current opinion: What is a nanoplastic?, Environ Pollut 235 (2018) 1030–1034. 10.1016/j.envpol.2018.01.024.

[3] S.L. Wright, F.J. Kelly, Plastic and Human Health: A Micro Issue?, Environ. Sci. Technol. 51 (2017) 6634–6647. 10.1021/acs.est.7b00423.

[4] P. Wick, A. Malek, P. Manser, D. Meili, X. Maeder-Althaus, L. Diener, P.-A. Diener, A. Zisch, H.F. Krug, U. von Mandach, Barrier capacity of human placenta for nanosized materials, Environ Health Perspect 118 (2010) 432–436. 10.1289/ehp.0901200.

[5] A. Ragusa, A. Svelato, C. Santacroce, P. Catalano, V. Notarstefano, O. Carnevali, F. Papa, M.C.A. Rongioletti, F. Baiocco, S. Draghi, E. D’Amore, D. Rinaldo, M. Matta, E. Giorgini, Plasticenta: First evidence of microplastics in human placenta, Environment International 146 (2021) 106274. 10.1016/j.envint.2020.106274.

[6] M.A. Garcia, R. Liu, A. Nihart, E. El Hayek, E. Castillo, E.R. Barrozo, M.A. Suter, B. Bleske, J. Scott, K. Forsythe, J. Gonzalez-Estrella, K.M. Aagaard, M.J. Campen, Quantitation and identification of microplastics accumulation in human placental specimens using pyrolysis gas chromatography mass spectrometry, Toxicological Sciences 199 (2024) 81–88. 10.1093/toxsci/kfae021.

[7] S.B. Fournier, J.N. D’Errico, D.S. Adler, S. Kollontzi, M.J. Goedken, L. Fabris, E.J. Yurkow, P.A. Stapleton, Nanopolystyrene translocation and fetal deposition after acute lung exposure during late-stage pregnancy, Part Fibre Toxicol 17 (2020) 55. 10.1186/s12989-020-00385-9.

[8] E.A. Medley, M.J. Spratlen, B. Yan, J.B. Herbstman, M.A. Deyssenroth, A Systematic Review of the Placental Translocation of Micro- and Nanoplastics, Curr Envir Health Rpt 10 (2023) 99–111. 10.1007/s40572-023-00391-x.

[9] J.P. Huang, P.C.H. Hsieh, C.Y. Chen, T.Y. Wang, P.C. Chen, C.C. Liu, C.C. Chen, C.P. Chen, Nanoparticles can cross mouse placenta and induce trophoblast apoptosis, Placenta 36 (2015) 1433–1441. 10.1016/j.placenta.2015.10.007.

[10] P. Wick, A. Malek, P. Manser, D. Meili, X. Maeder-Althaus, L. Diener, P.A. Diener, A. Zisch, H.F. Krug, U. Von Mandach, Barrier capacity of human placenta for nanosized materials, Environmental Health Perspectives 118 (2010) 432–436. 10.1289/ehp.0901200.

[11] Z. Yang, G.M. DeLoid, H. Zarbl, J. Baw, P. Demokritou, Micro- and nanoplastics (MNPs) and their potential toxicological outcomes: State of science, knowledge gaps and research needs, NanoImpact 32 (2023) 100481. 10.1016/j.impact.2023.100481.

[12] Z. Aghaei, J.G. Sled, J.C. Kingdom, A.A. Baschat, P.A. Helm, K.J. Jobst, L.S. Cahill, Maternal Exposure to Polystyrene Micro- and Nanoplastics Causes Fetal Growth Restriction in Mice, Environ. Sci. Technol. Lett. 9 (2022) 426–430. 10.1021/acs.estlett.2c00186.

[13] S. Wan, X. Wang, W. Chen, Z. Xu, J. Zhao, W. Huang, M. Wang, H. Zhang, Polystyrene Nanoplastics Activate Autophagy and Suppress Trophoblast Cell Migration/Invasion and Migrasome Formation to Induce Miscarriage, ACS Nano 18 (2024) 3733–3751. 10.1021/acsnano.3c11734.

[14] F. Amereh, N. Amjadi, A. Mohseni-Bandpei, S. Isazadeh, Y. Mehrabi, A. Eslami, Z. Naeiji, M. Rafiee, Placental plastics in young women from general population correlate with reduced foetal growth in IUGR pregnancies, Environmental Pollution 314 (2022) 120174. 10.1016/j.envpol.2022.120174.

[15] Y. Shen, Y. Ning, F. Chen, X. Wang, D. Zheng, Y. Sun, X. Zhang, X. Zhang, Impact of placental microplastics on birth anthropometrics: A cross-sectional study, Ecotoxicology and Environmental Safety 309 (2026) 119572. 10.1016/j.ecoenv.2025.119572.

[16] M. Ali-Hassanzadeh, N. Arefinia, Z.-A.-S. Ghoreshi, H. Askarpour, H. Mashayekhi-Sardoo, The effects of exposure to microplastics on female reproductive health and pregnancy outcomes: A systematic review and meta-analysis, Reprod Toxicol 135 (2025) 108932. 10.1016/j.reprotox.2025.108932.

[17] J. Gingrich, E. Ticiani, A. Veiga-Lopez, Placenta Disrupted: Endocrine Disrupting Chemicals and Pregnancy, Trends in Endocrinology & Metabolism 31 (2020) 508–524. 10.1016/j.tem.2020.03.003.

[18] J. Lv, Q. He, Z. Yan, Y. Xie, Y. Wu, A. Li, Y. Zhang, J. Li, Z. Huang, Inhibitory Impact of Prenatal Exposure to Nano-Polystyrene Particles on the MAP2K6/p38 MAPK Axis Inducing Embryonic Developmental Abnormalities in Mice, Toxics 12 (2024) 370. 10.3390/toxics12050370.

[19] N. Magosso, P.V. Souza, M.F. Moreira, V.A. Rocha, M.N. Fioretto, V.C. Pinha, G.A. Maia, V.L.R.S. Maria, L.A. Barata, G.F. Frigoli, G.S.A. Fernandes, A.C. Arena, W.R. Scarano, Maternal exposure to phthalates and nanoplastics, isolated or combined: Impacts on placental structure, development, and antioxidant defense as a trigger for maternal-fetal adversities, Reproductive Toxicology 135 (2025) 108930. 10.1016/j.reprotox.2025.108930.

[20] Z. Chen, M. Zheng, T. Wan, J. Li, X. Yuan, L. Qin, L. Zhang, T. Hou, C. Liu, R. Li, Gestational exposure to nanoplastics disrupts fetal development by promoting the placental aging via ferroptosis of syncytiotrophoblast, Environment International 197 (2025) 109361. 10.1016/j.envint.2025.109361.

[21] Y. Cheng, Y. Li, Y. Zhang, H. Liu, B. Yang, J. Zhu, H. Kuang, Gestational exposure to micro- and nanoplastics leads to poor pregnancy outcomes by impairing placental trophoblast syncytialization, Environmental Pollution 381 (2025) 126520. 10.1016/j.envpol.2025.126520.

[22] M. Nacka-Aleksić, A. Vilotić, A. Pirković, M. Živanović, B. Ljujić, M. Jovanović Krivokuća, Nano-scale dangers: Unravelling the impact of nanoplastics on human trophoblast invasion, Chemico-Biological Interactions 405 (2025) 111317. 10.1016/j.cbi.2024.111317.

[23] A. Ragusa, L. Cristiano, P. Di Vinci, G. Familiari, S.A. Nottola, G. Macchiarelli, A. Svelato, C. De Luca, D. Rinaldo, I. Neri, F. Facchinetti, Artificial plasticenta: how polystyrene nanoplastics affect in-vitro cultured human trophoblast cells, Front. Cell Dev. Biol. 13 (2025). 10.3389/fcell.2025.1539600.

[24] M. Horvat Mercnik, C. Schliefsteiner, G. Sanchez-Duffhues, C. Wadsack, TGFβ signalling: a nexus between inflammation, placental health and preeclampsia throughout pregnancy, Human Reproduction Update 30 (2024) 442–471. 10.1093/humupd/dmae007.

[25] K. Hashimoto, Y. Miyagawa, S. Watanabe, K. Takasaki, M. Nishizawa, K. Yatsuki, Y. Takahashi, H. Kamata, C. Kihira, H. Hiraike, Y. Sasamori, K. Kido, E. Ryo, K. Nagasaka, The TGF-β/UCHL5/Smad2 Axis Contributes to the Pathogenesis of Placenta Accreta, Int J Mol Sci 24 (2023) 13706. 10.3390/ijms241813706.

[26] Y. Yinon, O. Nevo, J. Xu, A. Many, A. Rolfo, T. Todros, M. Post, I. Caniggia, Severe Intrauterine Growth Restriction Pregnancies Have Increased Placental Endoglin Levels, Am J Pathol 172 (2008) 77–85. 10.2353/ajpath.2008.070640.

[27] S. Haider, A.I. Lackner, B. Dietrich, V. Kunihs, P. Haslinger, G. Meinhardt, T. Maxian, L. Saleh, C. Fiala, J. Pollheimer, P.A. Latos, M. Knöfler, Transforming growth factor-β signaling governs the differentiation program of extravillous trophoblasts in the developing human placenta, Proceedings of the National Academy of Sciences 119 (2022) e2120667119. 10.1073/pnas.2120667119.

[28] R. An, X. Wang, L. Yang, J. Zhang, N. Wang, F. Xu, Y. Hou, H. Zhang, L. Zhang, Polystyrene microplastics cause granulosa cells apoptosis and fibrosis in ovary through oxidative stress in rats, Toxicology 449 (2021) 152665. 10.1016/j.tox.2020.152665.

[29] K. Senathirajah, S. Attwood, G. Bhagwat, M. Carbery, S. Wilson, T. Palanisami, Estimation of the mass of microplastics ingested – A pivotal first step towards human health risk assessment, Journal of Hazardous Materials 404 (2021). 10.1016/j.jhazmat.2020.124004.

[30] N.H. Mohamed Nor, M. Kooi, N.J. Diepens, A.A. Koelmans, Lifetime Accumulation of Microplastic in Children and Adults, Environ. Sci. Technol. 55 (2021) 5084–5096. 10.1021/acs.est.0c07384.

[31] M. Pletz, Ingested microplastics: Do humans eat one credit card per week?, Journal of Hazardous Materials Letters 3 (2022) 100071. 10.1016/j.hazl.2022.100071.

[32] D. Xu, Y. Ma, C. Peng, Y. Gan, Y. Wang, Z. Chen, X. Han, Y. Chen, Differently surface-labeled polystyrene nanoplastics at an environmentally relevant concentration induced Crohn’s ileitis-like features via triggering intestinal epithelial cell necroptosis, Environment International 176 (2023) 107968. 10.1016/j.envint.2023.107968.

[33] A.B. Nair, S. Jacob, A simple practice guide for dose conversion between animals and human, J Basic Clin Pharm 7 (2016) 27–31. 10.4103/0976-0105.177703.

[34] C. Pan, X. Wang, Z. Fan, W. Mao, Y. Shi, Y. Wu, T. Liu, Z. Xu, H. Wang, H. Chen, Polystyrene microplastics facilitate renal fibrosis through accelerating tubular epithelial cell senescence, Food and Chemical Toxicology 191 (2024) 114888. 10.1016/j.fct.2024.114888.

[35] Q. Chen, C. Peng, R. Xie, H. Xu, Z. Su, G. Yilihan, X. Wei, S. Yang, Y. Shen, C. Ye, C. Jiang, Placental and fetal enrichment of microplastics from disposable paper cups: implications for metabolic and reproductive health during pregnancy, Journal of Hazardous Materials 478 (2024) 135527. 10.1016/j.jhazmat.2024.135527.

[36] S. Furukawa, Y. Kuroda, A. Sugiyama, A Comparison of the Histological Structure of the Placenta in Experimental Animals, Journal of Toxicologic Pathology 27 (2014) 11–18. 10.1293/TOX.2013-0060.

[37] M.W. Pfaffl, A new mathematical model for relative quantification in real-time RT–PCR, Nucleic Acids Res 29 (2001) e45.

[38] H. Alahmadi, M. Nadeem, A.M. Pujols, R. Reynolds, M.S. Islam, I. Gupta, C. Potts, A. Harbolic, G. Lafontant, S. Mitra, G.R. Warner, Polystyrene and polyethylene terephthalate nanoplastics differentially impact mouse ovarian follicle function, Environmental Pollution 386 (2025) 127228. 10.1016/j.envpol.2025.127228.

[39] A.J. Nihart, M.A. Garcia, E. El Hayek, R. Liu, M. Olewine, J.D. Kingston, E.F. Castillo, R.R. Gullapalli, T. Howard, B. Bleske, J. Scott, J. Gonzalez-Estrella, J.M. Gross, M. Spilde, N.L. Adolphi, D.F. Gallego, H.S. Jarrell, G. Dvorscak, M.E. Zuluaga-Ruiz, A.B. West, M.J. Campen, Bioaccumulation of microplastics in decedent human brains, Nat Med (2025) 1–6. 10.1038/s41591-024-03453-1.

[40] T. Braun, L. Ehrlich, W. Henrich, S. Koeppel, I. Lomako, P. Schwabl, B. Liebmann, Detection of microplastic in human placenta and meconium in a clinical setting, Pharmaceutics 13 (2021) 1–12. 10.3390/pharmaceutics13070921.

[41] H. Zhang, X. Ding, H. Zheng, Q. Ma, T. Zhang, Health Risks of Prenatal and Early-Life Microplastics Exposure: A Comprehensive Review, Environ. Health 4 (2026) 567–584. 10.1021/envhealth.5c00388.

[42] M. Xiao, Y. Yang, H. Alahmadi, A. Harbolic, G.M. Moreno, T. Yu, J. Liu, A. Guo, G.R. Warner, P.A. Stapleton, H. Chen, Rapid detection of microplastics and nanoplastics in seconds by mass spectrometry, Journal of Hazardous Materials 493 (2025) 138322. 10.1016/j.jhazmat.2025.138322.

[43] C.M. Cary, G.M. DeLoid, Z. Yang, D. Bitounis, M. Polunas, M.J. Goedken, B. Buckley, B. Cheatham, P.A. Stapleton, P. Demokritou, Ingested Polystyrene Nanospheres Translocate to Placenta and Fetal Tissues in Pregnant Rats: Potential Health Implications, Nanomaterials 13 (2023) 720. 10.3390/nano13040720.

[44] D. Yang, J. Zhu, X. Zhou, D. Pan, S. Nan, R. Yin, Q. Lei, N. Ma, H. Zhu, J. Chen, L. Han, M. Ding, Y. Ding, Polystyrene micro- and nano-particle coexposure injures fetal thalamus by inducing ROS-mediated cell apoptosis, Environment International 166 (2022) 107362. 10.1016/j.envint.2022.107362.

[45] J.M. Ward, S.A. Elmore, J.F. Foley, Pathology Methods for the Evaluation of Embryonic and Perinatal Developmental Defects and Lethality in Genetically Engineered Mice, Vet Pathol 49 (2012) 71–84. 10.1177/0300985811429811.

[46] Development of Structures and Transport Functions in the Mouse Placenta | Physiology | American Physiological Society, Physiology (n.d.). https://journals.physiology.org/doi/10.1152/physiol.00001.2005 (accessed March 22, 2026).

[47] S.A. Elmore, R.Z. Cochran, B. Bolon, B. Lubeck, B. Mahler, D. Sabio, J. Ward, Histology Atlas of the Developing Mouse Placenta, Toxicol Pathol 50 (2022) 60–117. 10.1177/01926233211042270.

[48] L. Woods, V. Perez-Garcia, J. Kieckbusch, X. Wang, F. DeMayo, F. Colucci, M. Hemberger, Decidualisation and placentation defects are a major cause of age-related reproductive decline, Nat Commun 8 (2017) 352. 10.1038/s41467-017-00308-x.

[49] A. Bhurke, J. Davila, J.A. Flaws, M.K. Bagchi, I.C. Bagchi, Exposure to di-isononyl phthalate during early pregnancy disrupts decidual angiogenesis and placental development in mice, Reproductive Toxicology 120 (2023) 108446. 10.1016/j.reprotox.2023.108446.

[50] B. Wen, H. Liao, W. Lin, Z. Li, X. Ma, Q. Xu, F. Yu, The Role of TGF-β during Pregnancy and Pregnancy Complications, Int J Mol Sci 24 (2023) 16882. 10.3390/ijms242316882.

[51] W.-J. Wu, C.-F. Lee, C.-H. Hsin, J.-Y. Du, T.-C. Hsu, T.-H. Lin, T.-Y. Yao, C.-H. Huang, Y.-J. Lee, TGF-β inhibits prolactin-induced expression of β-casein by a Smad3-dependent mechanism, Journal of Cellular Biochemistry 104 (2008) 1647–1659. 10.1002/jcb.21734.

[52] K.-Q. Zhao, H.-Y. Lin, C. Zhu, X. Yang, H. Wang, Maternal Smad3 deficiency compromises decidualization in mice, Journal of Cellular Biochemistry 113 (2012) 3266–3275. 10.1002/jcb.24204.

[53] H. Zhou, G. Fu, H. Yu, C. Peng, Transforming growth factor-beta inhibits aromatase gene transcription in human trophoblast cells via the Smad2 signaling pathway, Reprod Biol Endocrinol 7 (2009) 146. 10.1186/1477-7827-7-146.

[54] L.C. Smith, S. Moreno, L. Robertson, S. Robinson, K. Gant, A.J. Bryant, T. Sabo-Attwood, Transforming growth factor beta1 targets estrogen receptor signaling in bronchial epithelial cells, Respir Res 19 (2018) 160. 10.1186/s12931-018-0861-5.

[55] C.M. Cary, T.N. Seymore, D. Singh, K.N. Vayas, M.J. Goedken, S. Adams, M. Polunas, V.R. Sunil, D.L. Laskin, P. Demokritou, P.A. Stapleton, Single inhalation exposure to polyamide micro and nanoplastic particles impairs vascular dilation without generating pulmonary inflammation in virgin female Sprague Dawley rats, Particle and Fibre Toxicology 20 (2023) 16. 10.1186/s12989-023-00525-x.

